# Mouse Cardiac Regeneration Database (MCAREDB): a single-cell atlas of neonatal cardiac regeneration in mice

**DOI:** 10.64898/2025.12.22.696074

**Authors:** Anthony Shea, Jiayi Cui, Meiyi Li, Riley J. Leonard, Josh Bartz, Thanh Nguyen, Jianyi Zhang, Daniel J. Garry, Xiao Dong

## Abstract

**Background:** The adult mammalian heart exhibits minimal regenerative capacity, rendering most major cardiac injuries irreversible. In contrast, neonatal mice retain the ability to regenerate cardiac tissue for a brief period after birth. Although single-cell RNA sequencing (scRNA-seq) has deepened our understanding of this regenerative window, existing datasets remain dispersed across studies, hindering integrative and comparative analyses.

**Methods and Results:** To address this gap, we developed the Mouse CArdiac REgeneration DataBase (MCAREDB), a freely accessible, web-based resource that consolidates scRNA-seq datasets from multiple mouse cardiac regeneration studies. MCAREDB enables cell type-specific exploration of differential gene expression and pathway enrichment across key biological contexts, including neonatal versus postnatal stages and sham versus myocardial infarction surgeries. Core features include an interactive genome browser with epigenetic tracks, comparative gene expression visualization, pathway enrichment analysis, and bulk data download options.

**Conclusion:** By integrating diverse datasets into a unified and interactive framework, MCAREDB lowers technical barriers to data exploration and accelerates hypothesis generation in cardiac regeneration research. The resource is freely available at https://MCAREDB.org:3305.

## Background

Adult humans, along with other mammals, exhibit minimal cardiomyocyte renewal. For example, radiocarbon (^14^C) dating of cardiomyocyte DNA found fewer than half of all cardiac cells are replaced during the average lifespan of a human, and the annual rate of human cardiomyocyte renewal was just ∼0.5%.^1^ Also, mice and pig cardiomyocytes almost completely lose the proliferative capacity after the first week after birth.^2,3^ Thus, the heart ranks among the least regenerative tissues in adult mammals.^4,5^ Consequently, cardiac damage, such as myocardial infarction (MI), which results in the loss of around one billion cardiomyocytes, leads to irreversible fibrotic scarring rather than functional tissue repair.^4,6^ This fundamental limitation highlights a critical unmet need for regenerative therapies in cardiovascular medicine.

Unlike adult mammals, neonatal mice display remarkable cardiac regenerative capacity within their first week of life.^7,8^ As shown previously, one-day old (P1) mice can fully regenerate resected ventricular tissue, or re-muscularized the left ventricle after left anterior descending (LAD) coronary artery ligation^9^, within 21 days, restoring normal systolic function.^7^ This regenerative process is driven primarily by the proliferation of preexisting cardiomyocytes, involving sarcomere disassembly and dedifferentiation.^10,11^ Importantly, this regenerative state is temporally transient, diminishing sharply a week after birth (P7) and disappearing entirely by adulthood.^7,8^ Beyond this window, cardiac injury elicits a maladaptive response characterized by fibrotic scar formation and functional decline.^7,12^ This narrow temporal window makes neonatal mice an ideal model system for investigating a central question in regenerative biology: *Which molecular mechanisms and gene expression profiles enable cardiac regeneration in neonates, and what factors suppresses this ability in the postnatal period?*

Recent technological advances, particularly single-cell RNA sequencing (scRNA-seq), have provided powerful tools for investigating neonate cardiac injury response at high resolution. scRNA-seq has revolutionized cardiac regeneration research by enabling transcriptional profiling at the single-cell resolution across diverse developmental stages and injury responses.^13,14^ Unlike bulk transcriptomics, which masks tissue heterogeneity, scRNA-seq allows for the precise identification and characterization of distinct cell populations, including cardiomyocytes in transitional states,^15,16^ fibroblasts,^17^ endothelial cells,^18^ and immune populations.^19^ Revealing their contributions to the molecular landscape underlying regeneration and its subsequent silencing.^17,20,21^ However, transcriptional output is just one layer of cellular identity. Complementary multi-omics approaches, such as single-cell assay for transposase-accessible chromatin using sequencing (scATAC-seq) and chromatin immunoprecipitation sequencing (ChIP-seq), have thus become essential for capturing the upstream regulatory mechanisms.^17,22,23^ These data types map chromatin accessibility and histone modification landscapes, enabling researchers to link gene expression changes to specific regulatory elements and epigenetic remodeling events that govern regenerative potential.

Despite recent advances, there remains a gap in integrated, query friendly databases for cross-study comparisons and multi-omics analysis. Without unification, datasets remain siloed, impeding researcher’s ability to identify shared patterns, validate findings across studies, and generate novel hypotheses. To address this gap, we developed the **M**ouse **CA**rdiac **RE**generation **D**ata**B**ase (MCAREDB) to address this need. MCAREDB provides a centralized, web-based resource that integrates scRNA-seq data from multiple cardiac regeneration studies. The website hosts differential expression analyses, pathway enrichment results, and intuitive visualization tools, enabling researchers to explore cell-type specific gene regulation across developmental stages and injury conditions. By bridging isolated datasets and lowering technical barriers, MCAREDB acts as a platform to accelerate discovery and translation in cardiac regeneration research.

## Methods

### Database Architecture and Implementation

We adapted the database framework of the Human Cell Aging Transcriptome Atlas (HCATA) that we developed recently.^24^ In brief, all processed differential expression and pathway enrichment data are stored in *MySQL* (v8.0.42) and exposed via a stateless *REST API* (Node.js v18.19.1). The frontend is built with *Angular* (v20.1.2) and *Bootstrap* (v5.3), providing a user-friendly interface with built-in mobile device compatibility and accessibility features (**Figure 1A**).

**Figure 1.**
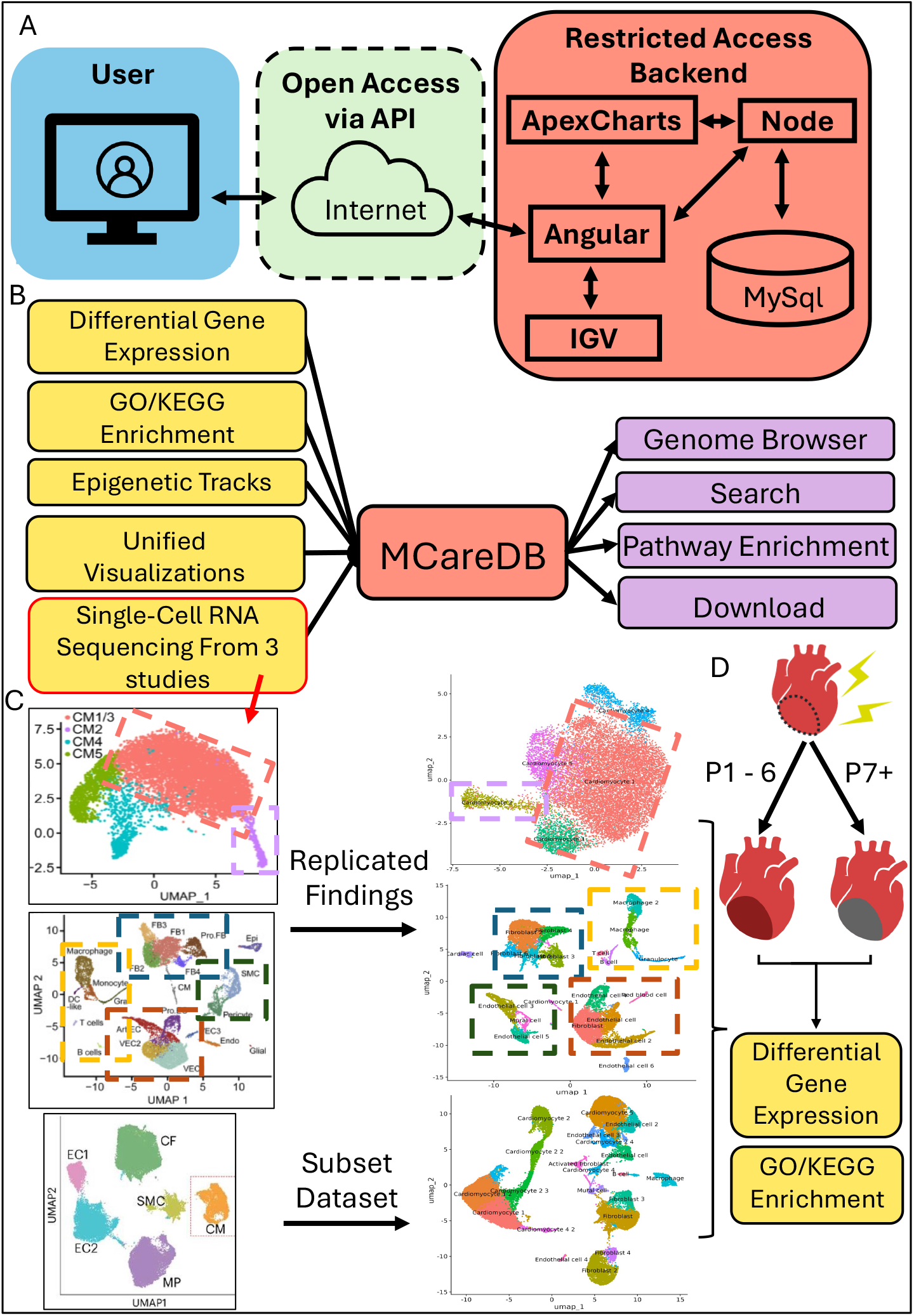
MCAREDB Architecture and Integrative Framework for Multi-Study Cardiac Regeneration Data. (A) Connections between the user interface, API, and backend components. (B) Data sources, analytical modules, and interactive elements that comprise MCAREDB. (C) Original UMAP from source publications and the MCAREDB counterpart UMAP. (D) Graphical representation of the regenerative capacity in neonatal (P1 - 6) and Postnatal (P7+) mice. Downstream analyses (Differential Gene Expression and Pathway Enrichment) are performed on both regenerative and non-regenerative hearts.

Website modules, including the Genome Browser, Gene Expression, Pathway Enrichment, and Search & Download page, retrieve precomputed data from the MySQL database. A server-side summary table stores precomputed statistics to optimize query performance. Visualization components utilize a customized fork of *IGV*.*js* (v3.4.0) for chromosomal rendering and *ApexCharts* (v4.7) for expression and pathway data display. The genome browser displays histone ChIP–seq signal (ENCFF657GDL; ENCSR675HDX) and whole-genome bisulfite-seq CpG coverage (ENCFF980ZXR; ENCSR397YEG) tracks obtained from the ENCODE Portal, both of which derive from 0-day old mice heart tissue (mm10).^25-28^

All frontend code and modifications to existing frameworks are published to a public GitHub repository (https://github.com/XiaoDongLab/Mouse-CArdiac-REgeneration-DataBase). Software tools and their versions used in conducting the database and website are provided in **Table S1**.

### Data Sources

We conducted a comprehensive literature search to identify datasets assessing cardiac regeneration in mice (**Table S2**). Datasets were retained if they met the following criteria: (i) comparisons between neonatal and postnatal stages, (ii) inclusion of myocardial infarction (MI) and sham surgery control groups, and (iii) consistent post-surgery sequencing dates (PSD1/3). Applying these criteria resulted in three datasets being retained for integration (**Figure 1B, C**).^17,21,29^

### Clustering and Cell-type Annotation

For each GEO series meeting our inclusion criteria, 10× matrix files were imported using *Seurat* (v4.4.0).^30^ Sample metadata, including natal phase, surgery type (Sham/MI), post-surgery sequencing day (PSD1/3), and genotype, were recorded. The processing workflow included data loading, quality control, normalization, integration, dimensionality reduction, clustering, and cell-type annotation. Cells were retained if they expressed more than 200 genes, fell below the 95^th^ percentile for gene counts, and contained less than 20% mitochondrial genes. Qualified cells were normalized using NormalizeData, and 2,000 highly variable genes were selected for downstream analysis. Data were integrated across samples using FindIntegrationAnchors (k = 50) and IntegrateData (k.weight = 30). Finally, Principal Component Analyses (PCA), Uniform Manifold Approximation and Projection (UMAP), and t-distributed stochastic neighbor embedding (t-SNE) were performed. Cardiomyocyte (CM) subpopulations were classified into five distinct clusters (CM1-5) using top differentially expressed marker genes derived from single-nuclei RNA sequencing (snRNA-seq) analysis. Additional marker genes were obtained from CellMarker 2.0.^31^ Clustering across the three studies yielded 14 major cell-types, comprising 28 distinct populations when including subtypes (**Table S3**).

### Differentially expressed genes (DEGs) and Pathway Enrichment Analysis

Based on the experimental design in Table 1 and Figure 1D, we performed two main comparisons: P1/P2 vs. P8 and MI vs. Sham, with the latter in each pair serving as the control. Differentially expressed genes (DEGs) were identified with associated p-values and log2 fold changes, and all genes were annotated by their Ensembl IDs.

**Table 1.** Experimental Groups.

| Main Comparison | Post-Surgery Sequencing Date | Sub-Group | Experimental Condition |
| --- | --- | --- | --- |
| Age: P1/P2 vs P8 | PSD1 | Sham | Sham – PSD1 |
|  |  | MI | MI – PSD1 |
|  | PSD3 | Sham | Sham – PSD3 |
|  |  | MI | MI – PSD3 |
| Surgery: Sham vs MI | PSD1 | P1/P2 | P1/P2 – PSD1 |
|  |  | P8 | P8 – PSD1 |
|  | PSD3 | P1/P2 | P1/P2 – PSD3 |
|  |  | P8 | P8 – PSD3 |

Differential expression was computed with FindMarkers with the Wilcoxon rank-sum test (min.pct = 0,1, logfc.threshold = 0). P-values were corrected for multiple testing using the Benjamini-Hochberg procedure. Gene symbols were mapped to Ensembl IDs via *org*.*Mm*.*eg*.*db* (v3.19).^32^ Ranked gene lists were subjected to Gene Ontology (GO) enrichment analysis^33^ using clusterProfiler^34^ (v4.10.0) and *fgsea* (v1.34.0, seed=123).^35^ Gene set sizes were constrained to [10, 500], and results with False Discovery Rate (FDR) < 0.05 and |Normalized Enrichment Score (NES)| > 1 were considered significant and retained for downstream interpretation. Due to out-of-date Kyoto Encyclopedia of Genes and Genomes^36^ (KEGG) pathway information in gseapy^37^, KEGG pathway enrichment analysis was performed using *rpy2* (v3.6) and *fgsea* (v1.34.0, seed=42) with KEGG pathway data. Same significance criteria (FDR < 0.05, |NES| > 1) were applied to KEGG enrichment results. These precomputed results, spanning all cell-types and experimental conditions, form the core analytical content of MCAREDB.

## Results

### Database Overview

MCAREDB is a centralized database used to store processed differential gene expression data and pathway enrichment results from mouse cardiac regeneration studies. Here we leverage original publications and associated metadata from three GEO series, comprising 20 samples that met our inclusion criteria (**Table S3**). Based on our data filtration and processing, we imported more than 2 million differential gene expression records, 1.7 million GO enrichment records, and more than 330,000 KEGG enrichment records. Each study contains scRNA-seq data from cardiac tissue of mice that: (1) underwent either MI or Sham surgery, (2) were sequenced 1 or 3 days after surgery, and (3) were either neonatal (0–7 days after birth) or postnatal (8+ days after birth). To minimize batch effects, studies were processed independently and are only displayed together in the database. Following cluster identification in dimensionality reduction plots, the top three most confident cell-type assignments were made based on gene expression profiles.

To identify differentially expressed genes (DEGs) associated with cardiac regeneration capacity, we performed pairwise comparisons between neonatal (P1) versus postnatal (P8) stages, myocardial infarction (MI) versus sham surgery controls, and sequencing date post-surgery (PSD1/3) across 20 samples from three GEO series (**Table S3**). Following quality control and normalization in Seurat, samples were integrated to minimize batch effects while preserving biological variation. Dimensionality reduction via PCA, UMAP, and t-SNE revealed 14 distinct cardiac cell populations, with cell-type identities assigned based on canonical marker gene expression. The raw data from each study were reanalyzed and clustered following the parameters of the source datasets. The results of the re-clustering confirm the consistency between the original publication and our reanalysis (**Figure 1C**). Clustering gives a clear separation of cardiomyocytes, fibroblasts, endothelial cells and immune cells which are usually presented in cardiac tissues, although the distribution pattern differs between our results and the original image due to potential parameter differences and clustering algorithms such as Leiden or Louvain (**Figure 1C**).^38,39^

### Exploratory analysis

Within each cell cluster, DEGs were identified using the Wilcoxon rank-sum test with Benjamini-Hochberg multiple testing correction. Genes showing differential expression (adjusted p-value < 0.05) between comparison groups were ranked by log2 fold change. Following DEG identification, we conducted gene set enrichment analysis using clusterProfiler and fgsea to identify enriched Gene Ontology (GO) biological processes and KEGG pathways.^34,35^ Both upregulated and downregulated gene sets were analyzed independently, with pathways meeting significance thresholds (FDR < 0.05, |NES| > 1) retained for downstream interpretation. These precomputed results, spanning all cell-types and experimental conditions, form the core analytical content of MCAREDB.

To explore the divergent repair profiles between neonatal and postnatal mice, we performed functional enrichment analysis using the R package clusterProfiler. Based on the identified DEGs, we present the top three most significantly enriched GO and KEGG terms across various relevant cell-types (**Figures 2 and S1**). GSEA was run separately for the two major comparison groups, Sham vs MI and P1 vs P8, to best capture what pathways are truly driving this regenerative capacity.

**Figure 2.**
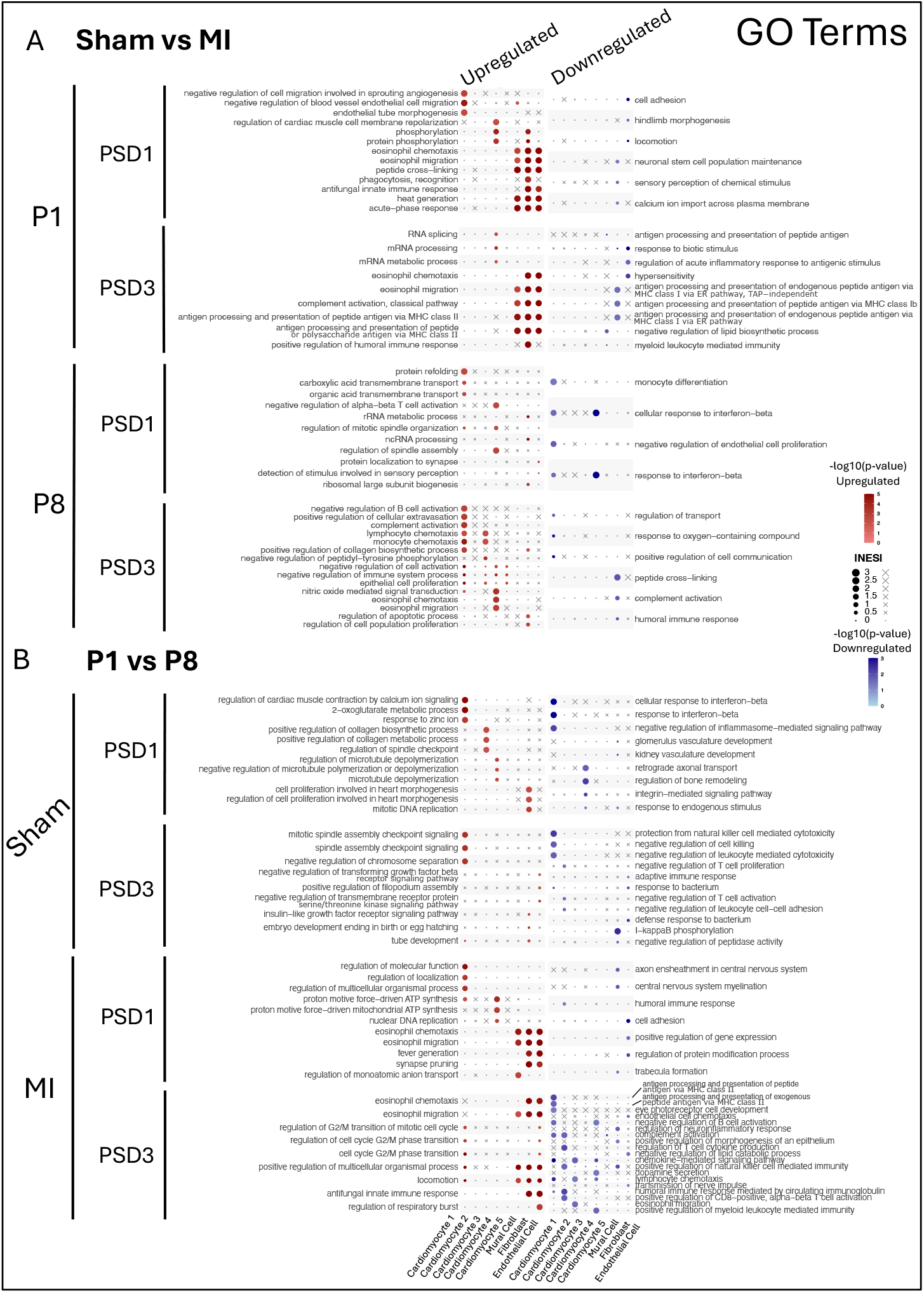
GO Pathway Enrichment Analysis between Age and Surgery Groups. (A) GO pathway enrichment between the Sham, control group, and Myocardial Infarction (MI), experimental group. (B) GO pathway enrichment between the P8, control group, and P1, experimental group.

To examine injury-specific response across developmental stages, GO pathway enrichment was performed on the Sham versus MI comparison (**Figure 2A**). At P1 PSD1 (1-day post-surgery), the cardiomyocyte 1 population upregulated a paradoxical anti-angiogenic response, while the cardiomyocyte 4 population displayed phosphorylation and cardiac muscle cell membrane repolarization, indicating active vascular regulation and electrophysiological remodeling (**Figure 2A**). Non-cardiomyocyte cells, including fibroblasts, mural cells, and endothelial cells, mounted robust acute-phase and eosinophil recruitment. This pattern is consistent with early muscle injury responses, where recruited eosinophils release IL-4 to activate muscle-resident progenitors and initiate regeneration.^40,41^ By P1 PSD3 (3-day post-surgery), the Sham versus MI comparison revealed a transition toward transcriptional reprogramming and a continuation of eosinophil recruitment. The cardiomyocyte 4 population now exhibited RNA processing pathways, aligning with previous characterizations of this population during cardiac damage showing enhanced ribosome biogenesis and protein synthesis.^21^ Non-cardiomyocyte cells maintained robust eosinophil recruitment, indicating sustained inflammatory cell recruitment during early repair. Surprisingly, all non-cardiomyocyte cells (Mural cells, fibroblasts and endothelial cells) showed upregulation of MHC Class II antigen processing and presentation in regenerative hearts (**Figure 2A**), whereas this response had previous only been demonstrated in Antigen Presenting Cells and fibroblasts for T cell activation.^42-44^

To complement the above findings in the injury specific response and to potentially isolate pathways specific to the neonatal heart regeneration, GO pathway enrichment was performed for P1 versus P8 across both surgery conditions (**Figure 2B**). Several cell-types demonstrated a robust regenerative signature characterized by active proliferation and developmental processes unique to the regenerative heart. In the P1 PSD1 (1-day post-MI) sample, cardiomyocytes showed an upregulation of broad regulatory and metabolic processes, with Cardiomyocyte 1 demonstrating increased activity across multiple regulatory pathways including molecular function, localization, and multicellular organismal processes, whereas Cardiomyocyte 4 populations characteristically showed enhanced mitochondrial ATP synthesis driven by proton motive force, along with modest nuclear DNA replication, indicating active energy production and some proliferative signaling.^21^ As previously shown in the Sham versus MI comparison, Non-cardiomyocyte populations (mural cells, fibroblasts, and endothelial cells) mounted eosinophil recruitment. Notably, P1 hearts showed minimal pathway suppression, with only sparse downregulation in select cardiomyocyte subtypes and stromal cells, indicating a predominantly upregulated profile that supports early regenerative mechanisms without substantial inhibitory signaling. On 3 days post-MI, the P1 neonatal cardiac injury response underwent substantial reprogramming characterized by focused cell cycle activation when compared to the P8 mice (**Figure 2B**). Cardiomyocyte 1 displayed upregulation of cell cycle regulatory processes governing the G2/M transition, alongside enhanced multicellular organismal regulation and locomotion, suggesting coordinated proliferative and migratory activity. Non-cardiomyocyte cells maintained robust eosinophil recruitment in addition to increased cellular coordination and locomotion.

### Web Interface Implementation

To support broad accessibility, we developed a user-friendly web interface that provides intuitive navigation of genomic and transcriptomic data. Core modules include an Interactive Genome Viewer for locus-level exploration, Gene Expression plots for comparing experimental conditions, a Pathway Enrichment page for investigating functional trends, and a Search & Download page for bulk downloads of all described analysis results. Together, these features allow for a friendly user interface to navigate between genome-wide summaries and cell-type–specific details.

The **Home** page of the website provides users with a brief description of the website as well as a brief *User Guide* to explain all the various components, including how the experiments of the original research was performed.

The **Genome Browser** page features an interactive genome browser for querying specific genes. This tool renders chromosomal loci alongside histone ChIP-seq (heart tissue, postnatal 0-day; doi:10.17989/ENCSR675HDX) and whole-genome bisulfite-seq (heart tissue, postnatal 0-day; doi:10.17989/ENCSR397YEG) tracks, enabling navigation from genome-level views down to individual genes (**Figure 3A**).^28^ For each queried gene, differential expression data is displayed across eight volcano plots, corresponding to each experimental condition (**Figures 3B-C** and **Table 1**). This module facilitates the comparison of transcriptomic differences in mouse hearts between the neonatal (P1-P6) and postnatal (P7+) phases, under Sham or MI conditions, and for 1- and 3-day PSD sequencing. To clarify the data, we defined genes with | log_2_ *FC* | > 0.58 (∼1.5 fold) as significantly up- or down-regulated, and those with 0.25 ≤ |log_2_ *FC* | < 0.58 (∼1.2-1.5 fold) were considered as slightly up- or down-regulated genes (**Table 2**).

**Table 2.**
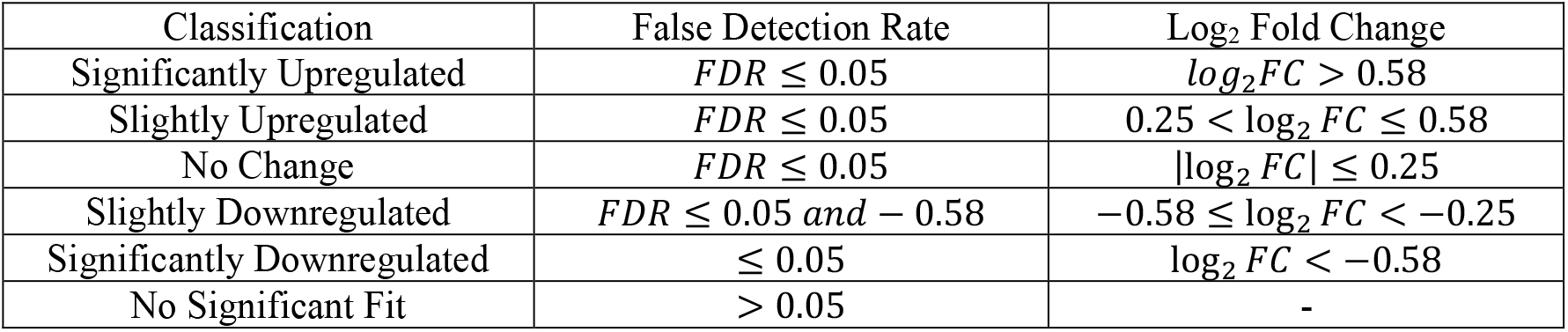
Differential Gene Expression Classification.

**Figure 3.**
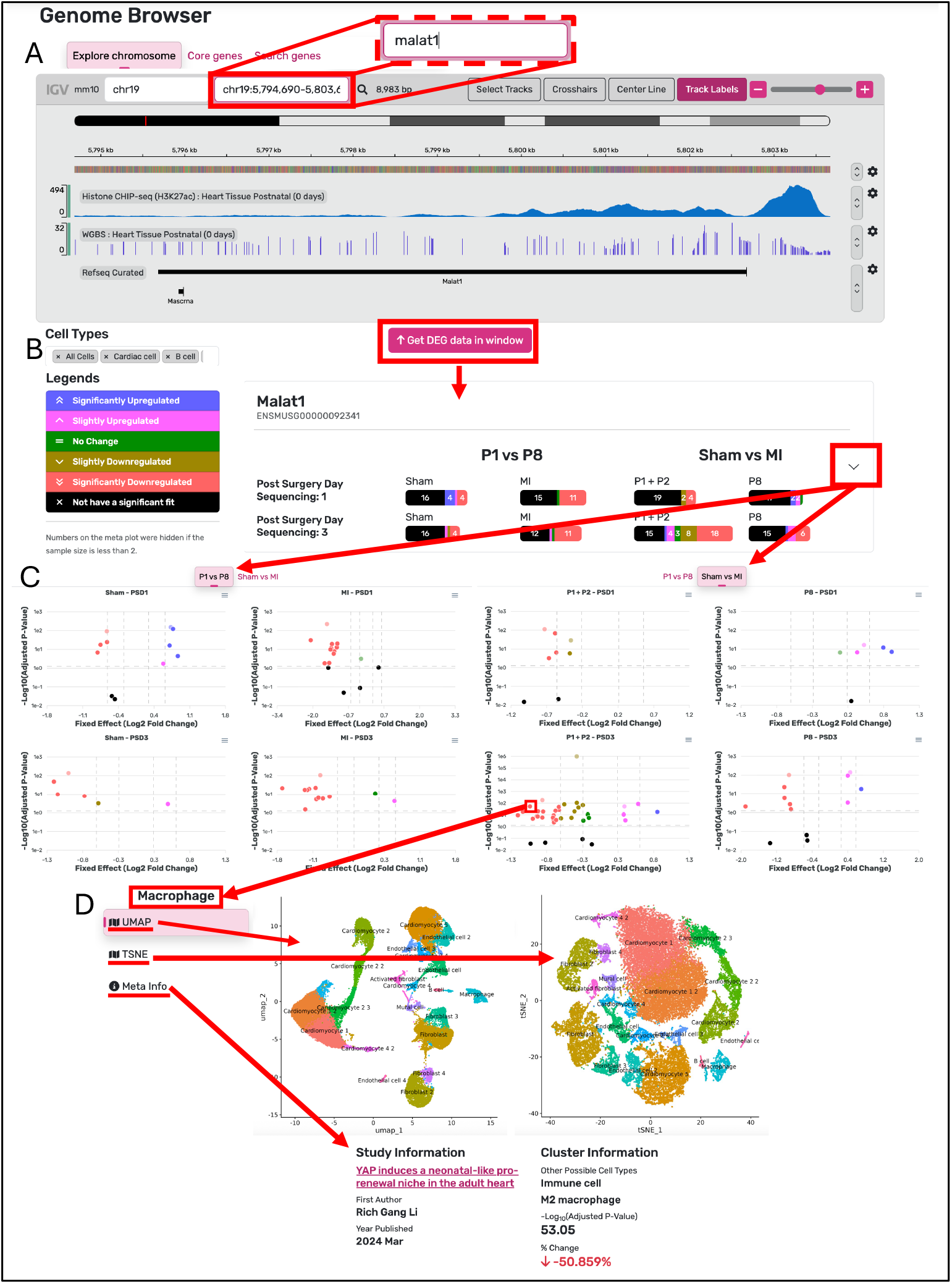
Interactive Genome Browser Interface. (A) Genome browser view showing gene and epigenetic features for MALAT1 in postnatal (0 days) heart tissue. (B) The fetch button retrieves all genes within the visible genome window and displays gene-specific results across comparison groups. (C) Expanding a gene via the dropdown arrow reveals volcano plots for each comparison group, highlighting cell-type–specific differential expression. (D) Selecting a point on a volcano plot opens study-specific details, including UMAP and t-SNE visualizations, study metadata, and cluster-level information.

Users can select any data point, which represents a cell-type, to view its corresponding UMAP, t-SNE, and statistical data (**Figure 3D**). Furthermore, the interface supports querying single genes, multiple genes at once, or a pre-defined list of prenatal heart regeneration genes for bulk analysis.

The **Gene Expression** page enables side-by-side comparison of genes across experimental groups (**Figure 4A**). Users can query a gene and then filter the results by Cell Type, Surgery Type, and Age Group. Each query adds a new data series to a centralized bar plot. To enhance clarity, several display options are available, including: a log scale toggle, the ability to separate data by sequencing date, and controls to show or hide specific experimental groups. Users can also remove specific genes from the plot using the “Remove Genes” function.

**Figure 4.**
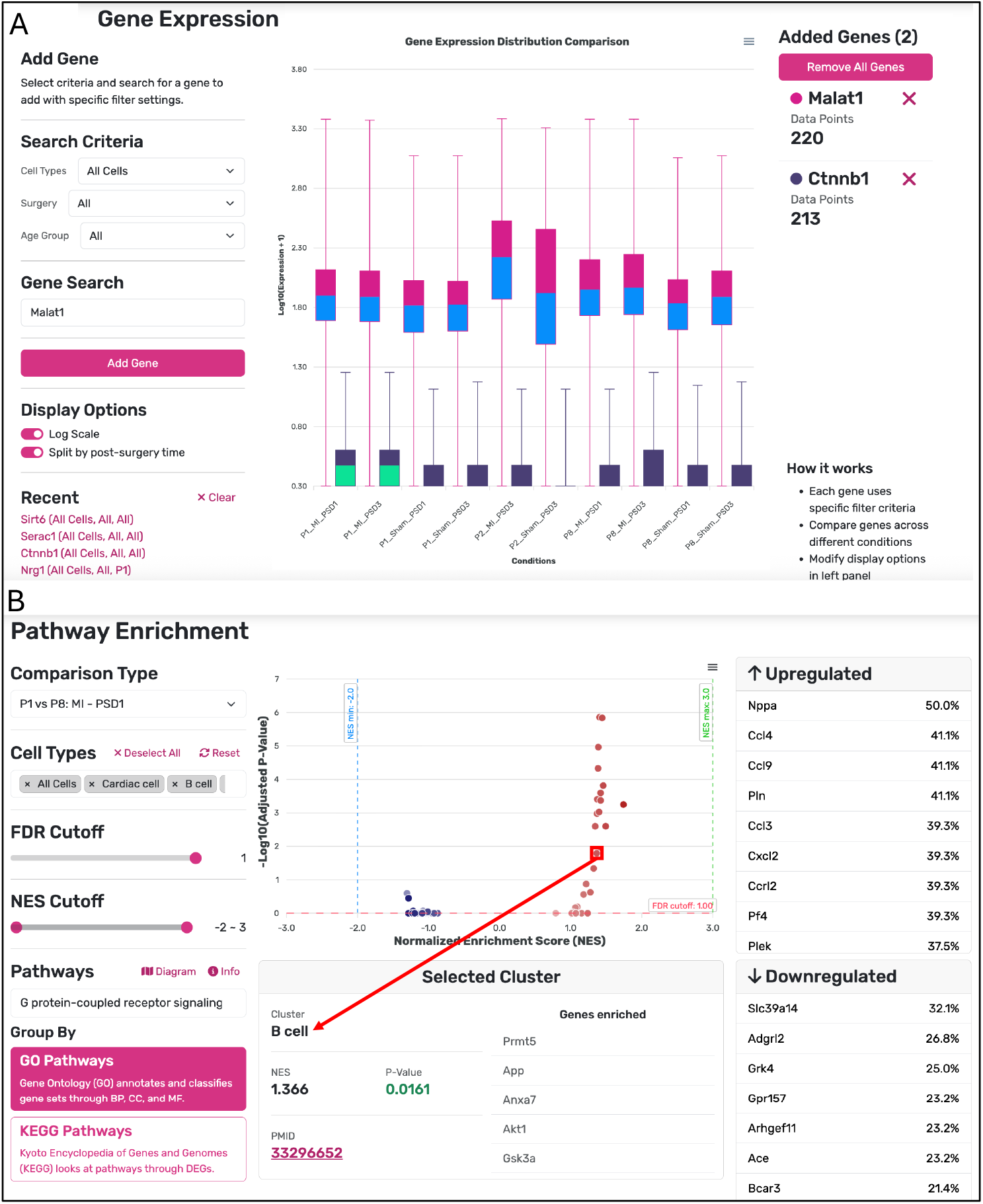
Gene Expression and Pathway Enrichment Analysis. (A) The Gene Expression tab displays queried expression values across comparison groups for example genes such as Malat1 and Ctnnb1. Users can adjust axes to log scale and choose to concatenate or separate data by surgery time groups. (B) The Pathway Enrichment tab shows a volcano plot of pathway enrichment results (Gene Ontology or Kyoto Encyclopedia of Genes and Genomes) for a selected comparison group. Clicking a cluster reveals cell-type–specific information and the genes enriched in that cluster. The upregulated and downregulated panels display the most frequently enriched genes, and users can filter results by false discovery rate (FDR) and normalized enrichment score (NES).

The **Pathway Enrichment** page enables users to explore differential gene expression within specific cell-types for GO and KEGG pathways under each experimental condition (**Figure 4B**). Users select either GO or KEGG to view the results on a central scatter plot, which displays the normalized enrichment score (NES) and statistical significance for each pathway. Clicking a data point opens a detailed panel containing meta-information for the cell-type, precise NES and FDR values, and a list of enriched genes. The page also lists the most common upregulated and downregulated genes across all clusters, sorted by prevalence. Selecting a gene navigates the user directly to its corresponding locus in the Genome Browser, creating a seamless connection from pathway-level analysis to genomic context.

The **Search & Download** page allows users to bulk download the analyses generated in this resource. All files can be filtered by File Type, Cell Type, and PMID.

The **Documentation** page provides definitions for all key terms used across the website, including Experimental Terminology, Statistical Terminology, and Cell type Definitions. For each cell-type identified on the site, the specific cell markers used to predict cluster identities are also listed.

## Conclusion

Accessible, integrated datasets for cardiac regeneration are lacking. MCAREDB addresses this gap by unifying disparate scRNA-seq studies into a cell-type resolved, queryable resource. Unlike general-purpose repositories, this database focuses specifically on neonatal versus postnatal cardiac regeneration, provides standardized data delivering visualization through a responsive web interface. By standardizing data processing and analysis, it removes technical barriers and accelerates the path from question to insight. Our comparative framework (Neonatal vs. Postnatal; Sham vs. MI) enables systematic identification of therapeutic targets and is designed to incorporate future datasets. Positioned at the interface of discovery and translation, MCAREDB establishes critical infrastructure to drive advances in cardiac regeneration research.

## Data and code availability

All data is available at the MCAREDB (https://MCAREDB.org:3305). Source code is available at GitHub (https://github.com/XiaoDongLab/Mouse-CArdiac-REgeneration-DataBase).

## Acknowledgement

This work was supported by the US National Institutes of Health (P01 HL160476, P01 AI172501, and U19 AG056278). The funders had no role in study design, data collection and analysis, decision to publish, or preparation of the manuscript. We appreciate the suggestions and comments from Zhenqing Ye at the University of Minnesota, Twin Cities.

---

**Figure S1.**
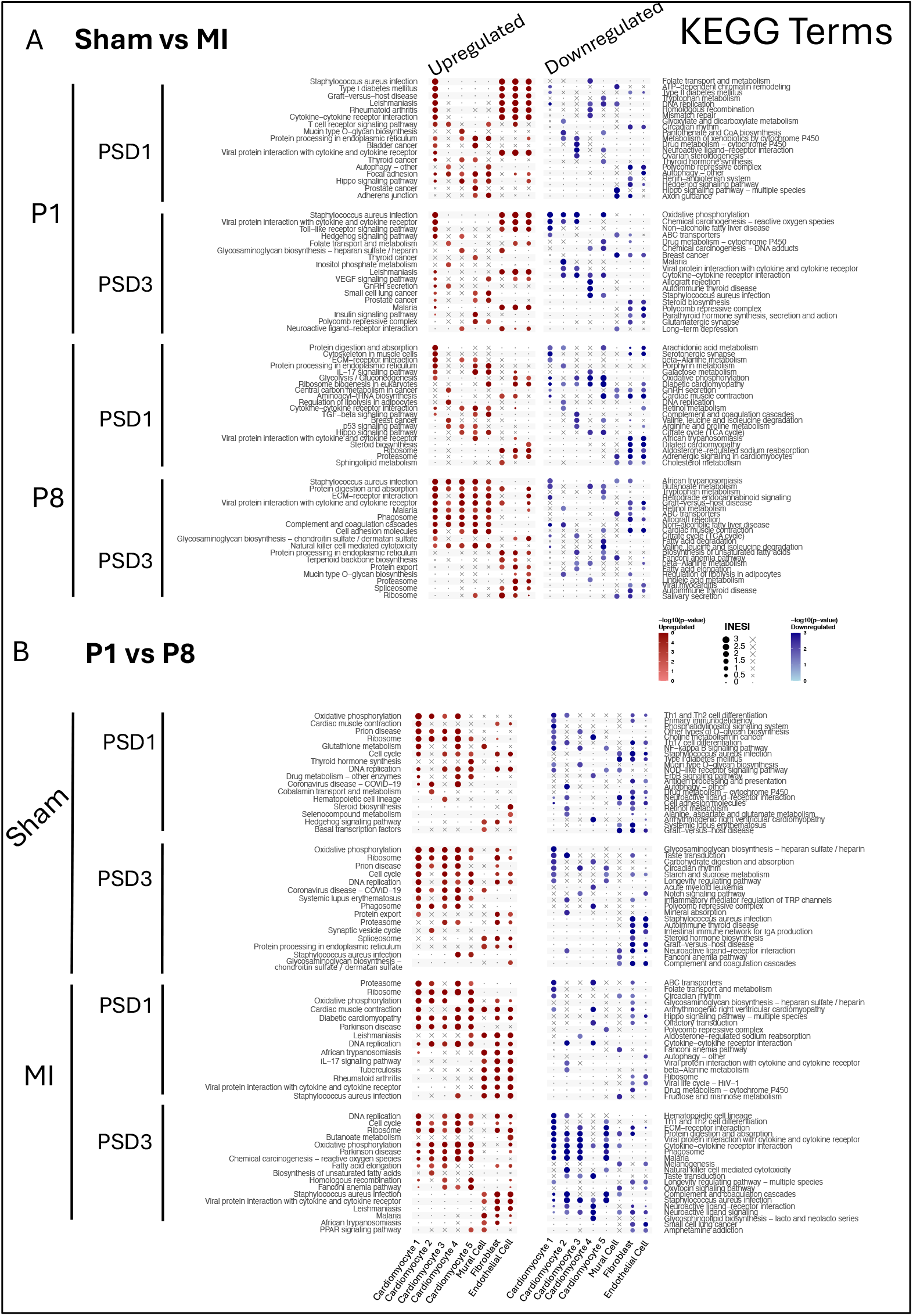
KEGG Pathway Enrichment Analysis between Age and Surgery Groups. (A) KEGG pathway enrichment between the Sham, control group, and Myocardial Infarction (MI), experimental group. (B) KEGG pathway enrichment between the P8, control group, and P1, experimental group.

**Table S1.** Software tools used in MCAREdb.

| Package | Version | GitHub Repository |
| --- | --- | --- |
| Angular | 20.1.2 | https://github.com/angular/angular |
| Node.js | 18.19.1 | https://github.com/nodejs/node |
| MySQL | 8.0.42 | https://github.com/mysql/mysql-server |
| Bootstrap | 5.3 | https://github.com/twbs/bootstrap |
| IGV.js<sup>45</sup> | 3.4.0 (modified) | https://github.com/Itsuki5696/igv-modern |
| ng-select | 15.0.0 | https://github.com/ng-select/ng-select |
| jszip | 3.10.1 | https://github.com/Stuk/jszip |
| Apexcharts | 4.7 | https://github.com/apexcharts/apexcharts.js |
| Ngx-translate | 16.0.1 | https://github.com/ngx-translate/core |
| Ag-grid | 33.3.2 | https://github.com/ag-grid/ag-grid |
| rxjs | 7.8.2 | https://github.com/ReactiveX/rxjs |
| Rpy2 | 3.6 | https://github.com/rpy2/rpy2 |

**Table S2.** Data source of MCAREDB.

| PMID | Sample ID | Age | Genotype | Surgery | Post Surgery Sequencing |
| --- | --- | --- | --- | --- | --- |
| 32220304 | GSM3747856 | P1 | ICR/CD1 | MI | 1 |
|  | GSM3747858 | P1 | ICR/CD1 | MI | 3 |
|  | GSM3747857 | P1 | ICR/CD1 | Sham | 1 |
|  | GSM3747859 | P1 | ICR/CD1 | Sham | 3 |
|  | GSM3747860 | P8 | ICR/CD1 | MI | 1 |
|  | GSM3747862 | P8 | ICR/CD1 | MI | 3 |
|  | GSM3747861 | P8 | ICR/CD1 | Sham | 1 |
|  | GSM3747863 | P8 | ICR/CD1 | Sham | 3 |
| 33296652 | GSM4644949_P1_1MI | P1 | ICR/CD1 | MI | 1 |
|  | GSM4644950_P1_1Sham | P1 | ICR/CD1 | Sham | 1 |
|  | GSM4644951_P1_3MI | P1 | ICR/CD1 | MI | 3 |
|  | GSM4644952_P1_3Sham | P1 | ICR/CD1 | Sham | 3 |
|  | GSM4644953_P8_1MI | P8 | ICR/CD1 | MI | 1 |
|  | GSM4644954_P8_1Sham | P8 | ICR/CD1 | Sham | 1 |
|  | GSM4644955_P8_3MI | P8 | ICR/CD1 | MI | 3 |
|  | GSM4644956_P8_3Sham | P8 | ICR/CD1 | Sham | 3 |
| 38510108 | GSM7658929_3 day post-MI | P2 | WT | MI | 3 |
|  | GSM7658930_3 day post-MI | P2 | WT | MI | 3 |
|  | GSM7658931_3 day post-Sham | P2 | WT | Sham | 3 |
|  | GSM7658932_3 day post-Sham | P2 | WT | Sham | 3 |

**Table S3.**
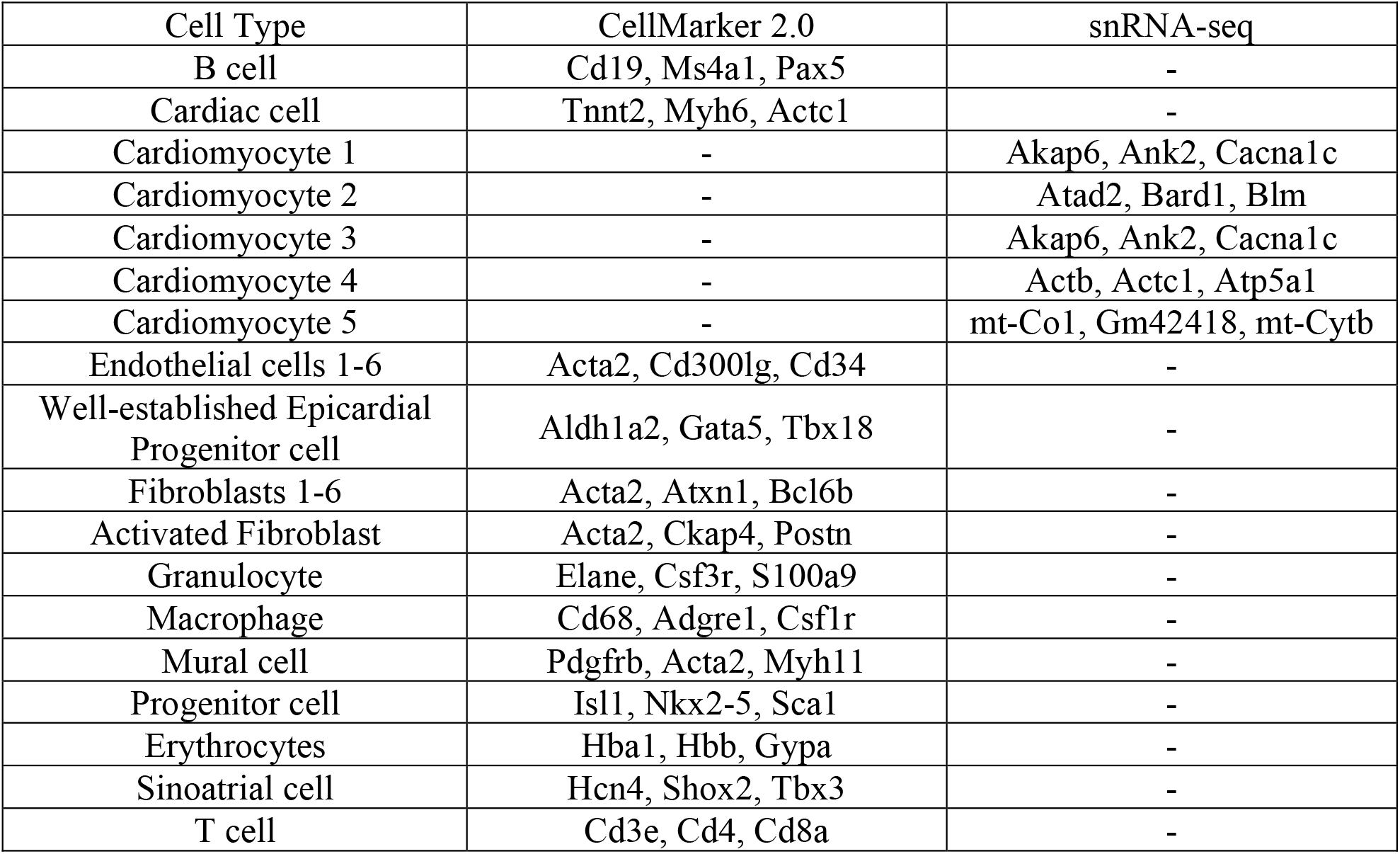
Gene Markers for Cell-Type Annotation.

